# Revision of the Community Conservation Index

**DOI:** 10.64898/2026.08.30.748128

**Authors:** Craig R. Macadam

## Abstract

The Community Conservation Index (CCI) developed by Chadd and Extence (2004) has been widely used to assess the conservation value of freshwater macroinvertebrate assemblages by integrating species rarity and community richness into a single metric. However, the original conservation scoring system was based largely on historical rarity designations and regional distribution data that no longer reflect current conservation assessments or recording coverage. This paper presents a comprehensive revision of the Conservation Scores underpinning the CCI. Species scores were updated using the latest IUCN-compliant Red List assessments for British freshwater invertebrates. For taxa classified as Least Concern, Data Deficient, or lacking a recent conservation assessment, scores were derived from national occurrence data across Great Britain for the period 1995-2024 using hectad occupancy. In addition, all non-native species were assigned a conservation score of zero to ensure they do not contribute positively to site conservation value.

The revised dataset includes 1,680 taxa, compared with 1,117 in the original index, with 755 taxa receiving Conservation Scores for the first time. Thirty non-native taxa were assigned a score of zero. Among previously scored taxa, 165 received higher scores, 576 received lower scores, and 184 remained unchanged. The predominance of score reductions reflects improved knowledge of species distributions and the replacement of historical rarity classifications with contemporary assessments based on conservation status and national frequency of taxon occupancy.

The revised CCI maintains the original strengths of combining rarity and richness while improving transparency, consistency and ecological relevance. By incorporating current Red List assessments, Great Britain-wide distribution data and revised treatment of non-native species, the updated index provides a more robust and nationally representative measure of freshwater macroinvertebrate conservation value. It offers an enhanced tool for biodiversity assessment, conservation prioritisation, site designation and environmental decision-making across freshwater ecosystems in Great Britain.

## Introduction

Freshwater macroinvertebrate communities are widely recognised as valuable indicators of ecological condition and conservation importance in rivers, streams, ponds and other inland waterbodies (Rosenberg and Resh, 1993). Their sensitivity to environmental change, coupled with extensive monitoring datasets, makes them a crucial component in assessing biodiversity, informing statutory protection, and guiding management interventions (Bonada et al., 2006). Traditional approaches to evaluating conservation value often relied primarily on the presence of rare or legally protected species, offering limited capacity to account for overall community richness or ecological representativeness (Pressey et al., 1993; Margules and Pressey, 2000). To address these limitations, Chadd and Extence (2004) developed the Community Conservation Index (CCI), a flexible, empirical tool that combines species rarity and community richness into a single measure of conservation value. Since its development, the Community Conservation Index has remained a widely used framework for assessing the conservation value of freshwater invertebrate assemblages and informing site-level conservation assessments (e.g. Hill et al., 2019; Kabir et al., 2023).

However, several aspects of the original scoring system now require revision in light of improved species distribution data, updated conservation assessments and modern biodiversity policy frameworks. First, the original conservation scores (CS) used in the CCI were based largely on the British Red Data Book (RDB) categories and National Rarity designations (Shirt, 1987; Bratton, 1991), many of which predate the adoption of internationally standardised IUCN Red List criteria (IUCN, 2012). These legacy classifications are not fully aligned with current threat assessment methodologies (Webb and Brown, 2016) and do not reflect recent changes in species status caused by climate change, habitat restoration, pollution mitigation or biological invasions (Strayer and Dudgeon, 2010; IPBES, 2023). In this revised version of the CCI, species-level conservation scores are instead derived from the most recent IUCN-compliant Red List assessments, ensuring that rarity scores are transparent, up to date and internationally comparable.

Second, the original lower-tier conservation scores (for common, frequent or local species) were informed primarily by distributional data from England. As national datasets have expanded and become more spatially complete (JNCC, 2025), this regional bias has become increasingly inappropriate. The updated CCI now assigns distribution-based scores using occurrence data from across the whole of Great Britain (England, Scotland and Wales). This provides a more accurate, nationally representative indication of frequency of taxon occupancy and avoids over- or under-valuing taxa that are locally scarce but widespread elsewhere.

Finally, the treatment of non-native species has been revised to reflect their lack of positive contribution to native biodiversity and, in many cases, their potential to negatively affect indigenous communities. In the original CCI, non-native or invasive taxa could inadvertently contribute to community richness without reducing the final index score. Under the revised system, all non-native species are assigned a conservation score of 0. This ensures they do not artificially inflate the conservation value of a site and recognises that their presence does not enhance, and may undermine, the ecological integrity of freshwater habitats (Feio et al., 2025).

Together, these refinements maintain the original purpose of the CCI - to provide a practical and scientifically robust measure of conservation value - while aligning it with contemporary conservation science, international criteria and broader geographic relevance. The updated index continues to balance species rarity and community richness but now provides a more accurate reflection of the conservation status of freshwater invertebrate assemblages across the Great Britain.

## Methods

Revision of the Conservation Scores was based on three components. First, conservation status, based on the IUCN Red List criteria (IUCN, 2012), for each species was sourced from the latest status reviews (Cook, 2015; Daguet et al., 2008; Foster, 2010; Macadam, 2015, 2016; Seddon, et al., 2014; Wallace, 2016). Scores were assigned according to Table 1.

**Table 1.** Revised Conservation Scores for freshwater taxa in Great Britain.

| CS | Definition |
| --- | --- |
| 10 | EX (Extinct) |
| 10 | CR (Critically Endangered) |
| 10 | CR(PE) (Critically Endangered Possibly Extinct) |
| 9 | EN (Endangered) |
| 8 | VU (Vulnerable) |
| 7 | NT (Near Threatened) |
| 6 | NR (Nationally Rare – found in less than 15 hectads) |
| 5 | NS (Nationally Scarce – found in less than 100 hectads) |
| 4 | Local (found in less than 291 hectads) |
| 3 | Frequent (found in less than 727 hectads) |
| 2 | Common (found in less than 1,455 hectads) |
| 1 | Very Common (found in 1,455 hectads or more) |
| 0 | Non-native species |

Where a species was classified as Least Concern or did not have a current status assessment, scores were based on the relative frequency within Great Britain using national occurrence data for the period 1995 to 2024, These data were sourced from the environment agencies (Environment Agency, Scottish Environment Protection Agency, and Natural Resources Wales) and combined with data from the NBN Atlas (www.nbnatlas.org.uk). Following Chadd and Extence (2004), species were placed in one of five categories based on the percentage of hectads (10 x 10 kilometres map squares) occupied across Great Britain (Table 1).

Data Deficient species were also scored using occupancy because insufficient evidence currently exists to assign a threat category, and distribution represents the most objective available measure of relative rarity.

Finally, all non-native species listed on the GB Non-Native Species Information Portal (https://www.nonnativespecies.org/non-native-species/information-portal), were assigned a score of 0. These species were retained in species lists but made no positive contribution to the conservation value.

## Results

The original CCI score covered 1,117 taxa. Since its publication, there have been a number of changes to nomenclature, and species have been synonymised, or species aggregations and complexes have been clarified with additional species being recognised. The new list of Conservation Scores comprises 1,680 taxa (Table 2; and Supplementary Information Table S1). 755 taxa were allocated a Conservation Score for the first time. Thirty non-native taxa were given a score of zero. Of those taxa that had previously been allocated a score, the Conservation Score was unchanged for 184 taxa.

**Table 2.** Number of taxa with changed Conservation Scores.

| Change | No. of Taxa |
| --- | --- |
| -5 | 3 |
| -4 | 42 |
| -3 | 85 |
| -2 | 218 |
| -1 | 228 |
| 0 | 184 |
| 1 | 125 |
| 2 | 25 |
| 3 | 11 |
| 4 | 3 |
| 5 | 1 |
| No previous score | 755 |
| Total | 1,680 |

165 taxa received a Conservation Score that was higher than the original score. The score for the majority of these taxa increased by a single point (Table 3). Only one taxon, the pea mussel *Euglesa ponderosa*, increased by five points.

**Table 3.** Breakdown down of Conservation Scores by Order.

| Order/Class | 0 | 1 | 2 | 3 | 4 | 5 | 6 | 7 | 8 | 9 | 10 | Total |
| --- | --- | --- | --- | --- | --- | --- | --- | --- | --- | --- | --- | --- |
| Acari |  |  |  |  |  |  | 1 |  |  |  |  | 1 |
| Annelida | 3 | 3 | 4 | 3 | 6 | 2 | 19 |  |  |  |  | 40 |
| Arachnida |  |  |  |  |  |  |  |  | 2 |  |  | 2 |
| Araneae |  |  |  | 1 |  |  |  |  |  |  |  | 1 |
| Bryozoa |  |  |  |  |  |  | 3 |  |  |  |  | 3 |
| Cnidaria |  |  |  |  |  |  | 1 |  |  |  |  | 1 |
| Coleoptera |  | 3 | 28 | 59 | 71 | 80 | 12 | 29 | 24 | 7 | 10 | 323 |
| Collembola |  |  |  |  | 1 | 3 |  |  |  |  |  | 4 |
| Crustacea | 15 | 2 |  | 3 | 7 | 26 | 13 |  |  |  |  | 66 |
| Diptera |  | 2 | 11 | 49 | 48 | 270 | 326 | 2 | 3 | 3 | 2 | 716 |
| Ephemeroptera |  | 1 | 14 | 8 | 10 | 8 | 5 |  | 2 | 3 | 2 | 53 |
| Hemiptera |  |  | 12 | 16 | 17 | 12 | 8 |  | 1 |  |  | 66 |
| Hymenoptera |  |  |  |  |  | 1 |  |  |  |  |  | 1 |
| Lepidoptera |  |  |  | 3 | 2 |  |  |  |  |  |  | 5 |
| Megaloptera |  |  | 1 | 1 | 1 |  |  |  |  |  |  | 3 |
| Mollusca | 9 | 2 | 12 | 15 | 9 | 18 | 13 | 1 | 8 | 1 | 3 | 91 |
| Neuroptera |  |  |  |  | 2 | 1 | 1 |  |  |  |  | 4 |
| Odonata |  | 13 | 8 | 5 | 5 | 2 | 2 | 6 | 2 | 4 | 3 | 50 |
| Plecoptera |  |  | 13 | 8 | 3 | 5 | 4 |  | 1 |  | 1 | 35 |
| Rotifera |  |  |  |  |  |  | 2 |  |  |  |  | 2 |
| Trichoptera |  | 8 | 27 | 43 | 28 | 47 | 27 | 5 | 8 | 3 | 4 | 200 |
| Turbellaria | 3 |  | 3 | 3 | 3 | 1 |  |  |  |  |  | 13 |
| **Total** | **30** | **34** | **133** | **217** | **213** | **476** | **437** | **43** | **51** | **21** | **25** | **1680** |

In contrast, 576 taxa received lower scores, with the majority decreasing by one or two points (Table 3). One taxon, the crane-fly *Tasiocera fuscescens*, received the largest change of five points. Two taxa, *Gammarus fossarum* and *Trocheta pseudodina*, which previously received a score of five, are now considered non-native species and were given a score of zero.

## Discussion

The revised Community Conservation Index (CCI) represents the most substantial update to the framework since its original publication by Chadd and Extence (2004). The expansion of the conservation score list from 1,117 to 1,680 taxa reflects both advances in taxonomic knowledge and the increased availability of biological recording data over the last two decades. This enlarged coverage improves the applicability of the index across a wider range of freshwater habitats and taxonomic groups, reducing the likelihood that ecologically important species are omitted from conservation assessments simply because they were absent from earlier scoring systems.

A key outcome of the revision is the widespread reduction in conservation scores among previously assessed taxa. Of the taxa reassessed, over three times as many experienced a decrease in score as an increase. This pattern is likely to reflect several factors. First, the original CCI relied heavily on historical rarity categories that were often based on incomplete distributional information. The substantial growth in biological recording effort (JNCC, 2025), improved digitisation of historical records, and the integration of national datasets have revealed many species to be more widespread than was previously understood. Consequently, some taxa formerly considered scarce can now be assigned lower conservation values that more accurately reflect their contemporary distribution. This does not imply that these species are less important biologically, but rather that the conservation scores are now based on a more robust evidence base.

The adoption of IUCN-compliant Red List assessments provides an important methodological improvement. Unlike earlier rarity-based approaches, IUCN assessments incorporate a broader range of criteria, including population trends, geographic range, habitat quality, and extinction risk (IUCN, 2012). By linking conservation scores directly to internationally recognised threat categories, the revised CCI better reflects genuine conservation concern rather than rarity alone. This is particularly important because rarity and vulnerability are not always closely correlated; some geographically restricted species may have stable populations, while widespread species can experience rapid declines (Gaston, 1994). The revised scoring framework therefore aligns more closely with contemporary conservation priorities and policy objectives.

The use of Great Britain-wide distribution data for assigning scores to taxa lacking a recognised threat status also addresses a significant limitation of the original index. Conservation assessments based solely on English distribution data may be geographically biased, as biodiversity data and conservation evidence are unevenly distributed across regions, and patterns identified in one country may not be representative of species’ responses elsewhere (Christie et al., 2020). By using a national distribution dataset, the revised approach provides a more representative measure of species frequency and facilitates more consistent comparisons between sites across Great Britain. This is especially relevant in the context of national conservation planning, where assessment tools should reflect biodiversity patterns at the scale at which policy and resource allocation decisions are often made (Margules and Pressey, 2000; Sarkar et al., 2006).

Although the use of hectad occupancy provides an objective and nationally consistent basis for assigning conservation scores, it is important to recognise that occupancy is a proxy for rarity rather than a direct measure of conservation status. Rarity is multidimensional, encompassing geographic range, habitat specificity and local abundance (Rabinowitz, 1981; Gaston, 1994). Consequently, species with restricted distributions but large local populations may receive relatively high conservation scores despite being numerically abundant, whereas widespread species with declining populations may receive lower scores than would be suggested by their conservation status alone. The incorporation of IUCN Red List assessments for threatened species helps partially address this limitation by ensuring that extinction risk is recognised where evidence is available. For species lacking formal assessments, however, occupancy provides the most practical and consistently available measure for application across the full range of freshwater invertebrate taxa considered by the CCI.

The decision to assign non-native species a conservation score of zero reflects an increasing emphasis on ecological integrity and the maintenance of native biodiversity within biodiversity assessments, consistent with approaches such as the Biodiversity Intactness Index that evaluates the condition of ecosystems based on persistence of originally occurring native species (Scholes and Biggs, 2005). Non-native taxa do not contribute to the conservation of indigenous biodiversity and some may negatively affect native freshwater communities through competition, predation, habitat alteration, or disease transmission (Feio et al., 2025). Under the revised system, their inclusion in species inventories continues to provide valuable ecological information while ensuring that they do not artificially increase site conservation scores. This change strengthens the ecological interpretation of the CCI and avoids potential inconsistencies whereby sites with high numbers of non-native species could previously receive elevated conservation values.

Despite these improvements, some limitations remain. Distribution-based measures of rarity are dependent on recording intensity and geographic coverage, which can vary substantially among taxonomic groups and regions. Although the use of data from 1995 to 2024 represents the most comprehensive evidence currently available, some species may still appear rarer or commoner than they truly are because of uneven survey effort (Isaac and Pocock, 2015). Similarly, Red List assessments are not updated uniformly across all freshwater invertebrate groups, meaning that some conservation scores may require revision as new assessments become available. Regular updates will therefore be necessary to maintain the scientific relevance of the index.

The revised CCI should be viewed as a dynamic framework rather than a fixed classification system. The increasing availability of biological records through national recording schemes, environmental agencies and the NBN Atlas provides opportunities for future refinements, including periodic recalibration of frequency thresholds and the incorporation of updated threat assessments. Such an adaptive approach will ensure that the index remains responsive to changes in species distributions, conservation status and environmental conditions.

In conclusion, the revised Community Conservation Index retains the strengths of the original methodology while addressing several important shortcomings. By incorporating contemporary Red List assessments, using Great Britain-wide distribution data, expanding taxonomic coverage, and excluding non-native species from contributing to conservation scores, the updated index provides a more robust and transparent assessment of freshwater invertebrate conservation value. As a result, it offers an improved tool for site evaluation, conservation prioritisation, monitoring and decision-making across freshwater ecosystems in Great Britain.

## Data Availability

The online version includes the full list of updated scores as supplementary information.

## Ethics and Permit Approval

No ethical approval was required.

## Use of Artificial Intelligence

The author used Microsoft Copilot to check the text and bibliography.

